# Revealing the structure of somatic cell membranes by *in situ* cryo-electron tomography

**DOI:** 10.1101/2022.09.19.508494

**Authors:** Chao Liu, Yiting Zhou, Tianyi Zou, Guanfang Zhao, Jinrui Zhang, Huili Wang, Hongda Wang

**Affiliations:** State Key Laboratory of Electroanalytical Chemistry, Changchun Institute of Applied Chemistry, Chinese Academy of Sciences, Changchun, Jilin 130022, China; University of Science and Technology of China, Hefei, Anhui 23006, China

## Abstract

The cell membrane, which separates the cell from the environment, plays a key role in signal transduction, energy conversion and substance transport. Although previous membrane models have successfully interpreted some functions of the cell membrane, no consensus has been reached for the lack of direct and *in situ* evidence. Here, we characterized the high-resolution 3D structure of 293T cell membranes *in situ* for the first time by cryo-electron tomography. Due to the excellent thickness of our cryo-samples, we could clearly observe membrane proteins with an average molecular weight of 100 kD. By analysing tomograms, we found that the total thickness of a 293T plasma membrane is approximately 20 nm and that there is a 4-nm lipid bilayer structure within the membrane. We observed that membrane proteins and protein complexes with a similar height (7-11 nm) are densely embedded in the ectoplasmic side of 293T plasma membranes, whereas membrane proteins aggregate to form islands with heights reaching dozens of nanometres on the cytoplasmic side. Additionally, we measured the average sizes of membrane proteins on the cytoplasmic side of 293T plasma membranes and found them to be approximately 7 nm in length and 4 nm in width. Moreover, if more precise structural information is obtained in future studies, we will identify the molecular interactions and detailed structures of membrane protein clusters that can be easily distinguished on a 293T cell membrane. Our work represents the first *in situ* structural characterization of a native somatic cell membrane with cryo-electron tomography and advances cell membrane structural studies from the model prediction stage to the real structure observation stage.

---

Cell membranes act as the first barrier to separate the cell from the environment and shield the cell from external harm. Consisting of thousands of types of lipids, proteins and saccharides, cell membranes are key elements for substance transport and the initiation of intracellular signals, among other functions(*1*). Although enormous varieties of membrane proteins, lipids and saccharides have been identified utilising proteomic and mass spectrometry analysis(*2, 3*), the relative positions, or rather the specific locations, of these fundamental components have mainly remained imprecise and contentious since the cell membranes were defined.

Great efforts have been made to demonstrate the structure of cell membranes from the beginning of the 20th century to the present. First published in 1925, Gorter and Grendel extracted the lipids with acetone and measured the lipid area by Langmuir methods, supposing that the membrane consists of a double lipid layer(*4*). A decade later, Danielli and Davson proposed a sandwich membrane model containing a very thin lipid layer covered by a protein layer on both sides(*5*). The suggestion that the protein layer was formed by direct physical adsorption provoked a spirited debate. Using electron microscopy, Sjostrand et al. observed a light band sandwiched between two dark bands after fixation and staining with heavy metals(*6*). Subsequently, the “unit membrane” model was proposed when Robertson identified the dark electron-dense bands as the lipid bilayer set with bilateral protein layers, bringing the concept of bilayer structure universally to all membrane systems(*7*). The Davson-Danielli model predominated until 1972, when Singer and Nicolson proposed the fluid mosaic model (FMM). As first depicted, the FMM represents plasma membranes as a matrix composed of a highly fluid bilayer of phospholipids with isolated membrane proteins as well as glycoproteins randomly intercalated into the plane of the membranes(*8*). Despite being the most representative general model of the plasma membrane, the FMM did not provide sufficient explanation of membrane functions, such as multiple protein-associated signal transduction and membrane endocytosis. Thus, Wang et al. proposed a protein layer–lipid–protein island (PLLPI) model that demonstrated the plasma membrane structure and functional mechanisms more precisely. As manifested in the model’s name, there is a dense protein layer (4 nm) over a lipid bilayer on the ectoplasmic side of the plasma membrane, while regularly dispersed protein aggregates form islands with a height of approximately 10–12 nm on the cytoplasmic side. The PLLPI model highlighted that the compact protein layer plays a vital role in the mechanical properties, signal transduction and substance transport of the membrane(*9*). However, ascertaining the accurate locations and relevant structures of membrane proteins at atomic resolution *in situ* remains a challenge for structural biologists(*10*).

Cryo-electron microscopy (cryo-EM) combines the high-resolution potential of 3D molecular-level imaging with the pristine structural preservation provided by vitrification by rapid freezing(*11*). Due to the development of high-resolution cryo-EM and the introduction of many new membrane mimetics, more than 1000 unique membrane protein structures have now been deposited. However, extraction from native membranes may change the structure and properties of membrane proteins, leading to conflicting results(*12*). Thus, it would be more authentic if the pure plasma membranes were observed *in situ* by cryo-EM. Although traditional methods, for example, sucrose gradient centrifugation and membrane patch unroofing procedures, have been utilised for plasma membrane preparation(*13, 14*), neither of them can mostly eliminate cellular content and fulfil the requirements of vitrification for cryo-ET, such as suitable thickness and homogenous fractions.

To achieve new structural and functional insights regarding plasma membranes, we obtained relatively clean plasma membrane cryo-samples directly on EM grids, which can be visualized *in situ* by cryo-EM. Taking advantage of cryo-electron tomography (cryo-ET) imaging, the overall distribution and specific locations of membrane proteins were observed with a high contrast on the extremely thin plasma membrane we prepared. Thus, more detailed and comprehensive plasma membrane structures of characteristic cells can be revealed by cryo-ET with the generalization of our plasma membrane preparation method.

## Distribution of membrane proteins observed with cryo-ET imaging

Owing to their superb transfection properties, we chose 293T cells to analyse the fundamental plasma membrane structure. Clean 293T cell membranes were freshly prepared directly on EM grids and then rapidly vitrified and loaded into a 300 kV Titan Krios cryo-electron microscope for cryo-ET imaging.

To accurately locate the area of plasma membranes by cryo-EM, we summarized empirical criteria to distinguish single-layer plasma membranes from thick cell samples. For example, in a magnified cryo-EM image of the 293T cell membrane, the opaque areas with a normal cell size were thought to be the intact 293T cells, and the almost transparent suborbiculate areas with a diameter ranging from 12 to 17 μm where we could clearly see the edge of holes on the supporting carbon film underneath were considered to be the single-layer 293T plasma membranes (Fig. 2, A1-A2). Through further tilt series acquisition from these membrane regions at higher magnification, more detailed structural features were observed in the correlative tomograms (Fig. 2, A3-A4).

To study the real morphology of intact and continuous plasma membranes, we collected data from relatively thin areas that were parts of recognizable cell membrane regions. The tilt series of our data were reconstructed into tomograms with IMOD. Intriguingly, membrane proteins with a relatively high contrast were clearly observed on an XY section of a representative plasma membrane tomogram (Fig. 1, B), and the profile of this plasma membrane can be easily determined from a relevant XZ section (Fig. 1, D). Additionally, we demonstrated the intuitive distribution of membrane proteins from the lateral (Fig. 1, C) and top views (Fig. 1, E) of a plasma membrane within its native environment by performing threshold segmentation and 3D visualization of its reconstruction results with Amira.

**Fig. 1.**
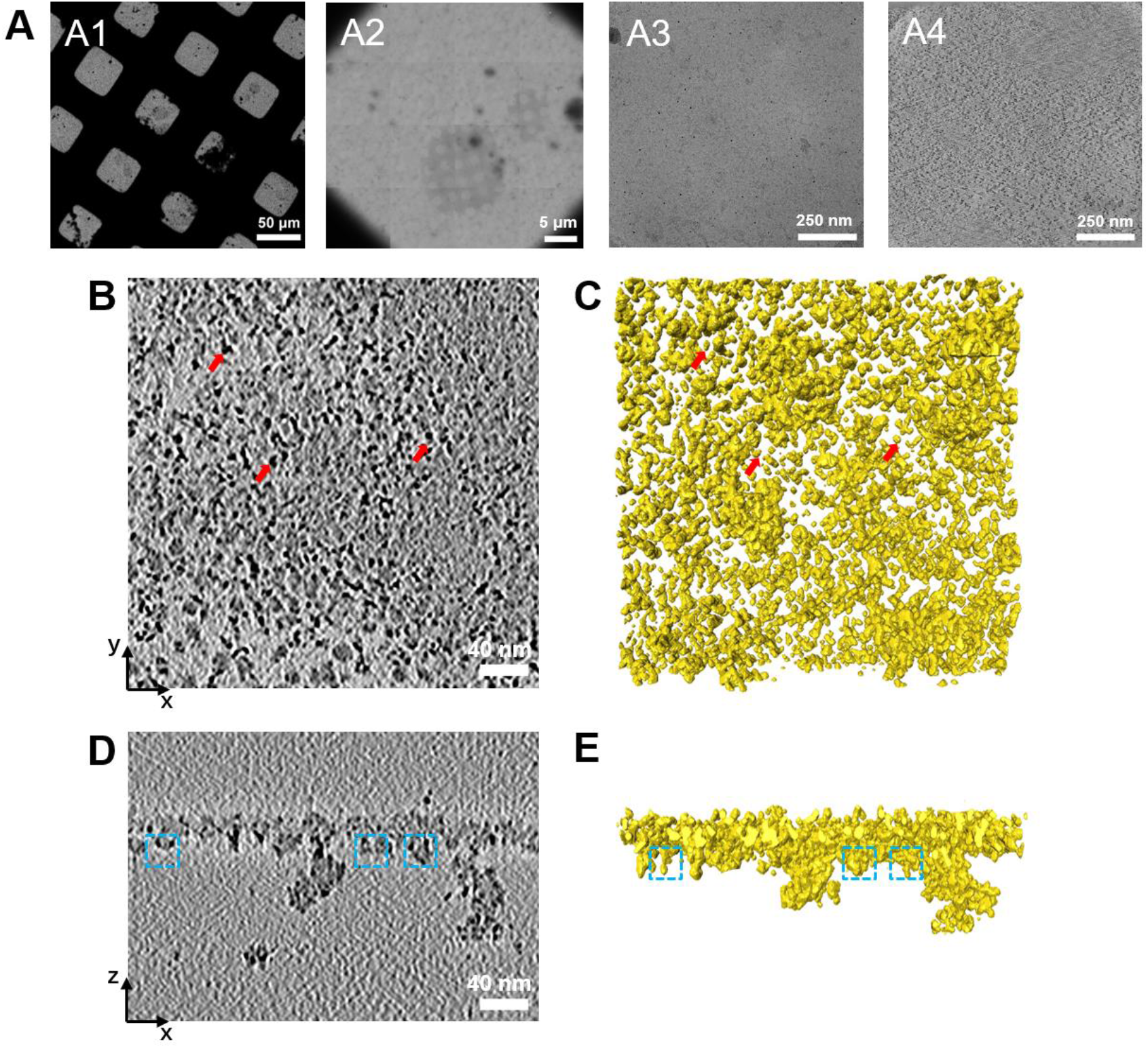
Cryo-ET Images and *in situ* structure of 293T plasma membrane. **(A)** Cryo-ET images of plasma membranes. **(A1)** A Cryo-ET image of recognizable cell contours on an EM grid at low magnification. **(A2)** A Cryo-ET image of membrane patches on one square of an EM gird. **(A3)** An image at zero angle in a tilt series showing indefinite features due to low contrast. **(A4)** A typical density slice from the 3D tomograms reconstructed from the tilt series of (A3) showing structural details. **(B)** An electron tomographic slice of a plasma membrane in a top view. **(C)** 3D rendering of the plasma membrane structure in the sub-tomogram shown in (B), the membrane proteins are pointed by red arrows. **(D)** An electron tomographic slice from a reconstructed sub-tomogram in a side view, showing the overall structure of a plasma membrane. The area above this bilayer structure is the ectoplasmic side of a plasma membrane, and the cytoplasmic area is under this bilayer. **(E)** 3D rendering of the plasma membrane structure in the sub-tomogram shown in (D). Membrane protein islands stand on the cytoplasmic side of a plasma membrane, which were marked by blue dashed boxes.

### Identification of the components in the native somatic plasma membranes

To determine the plasma membrane orientation in tomograms, the supporting side of an EM grid on which plasma membranes were deposited was always adjusted to face the electron beams in cryo-EM. In addition, the relative positions of Au nanoparticles that were dropped on the exposed cytoplasmic side of the plasma membrane provided another clue to confirm our judgement of the membrane orientation.

As the cytoplasmic and ectoplasmic side of a plasma membrane were determined, the whole membrane structure could be visualized by scanning sections of a tomogram generated from a tilt series of a membrane area along the z-axis (Fig. 2, A1-A4). At the far end of the ectoplasmic side of the membrane, there were quantities of large membrane protein complexes with relatively low contrast surrounded by dense membrane proteins with the same heights, providing a smooth surface (Fig. 2, A1). As depicted in images at high magnification, membrane complexes on the ectoplasmic side consisted of multiple membrane proteins that interact with each other, forming functional domains (Fig. 2, the magnified image of the area circled in red in the top right white box of A1). Switching sections from the ectoplasmic side to the cytoplasmic side, a highly continuous region with a thickness of 4 nm appeared on the ectoplasmic side of the membrane, where the membrane proteins were more crowded so that their signals became inadequate to distinguish them from each other (Fig. 2, A2). Progressing through slices along the z-axis, the signals of membrane proteins in sections with a thickness of 4 nm became easier to recognize due to the increasing gap between proteins. We speculated that these sections were a lipid bilayer with relatively less crowded but more identifiable transmembrane protein signals (Fig. 2, A3). For the cytoplasmic side of the membrane, membrane proteins were more likely to aggregate to form islands with heights reaching dozens of nanometres (Fig. 2, A4).

**Fig. 2.**
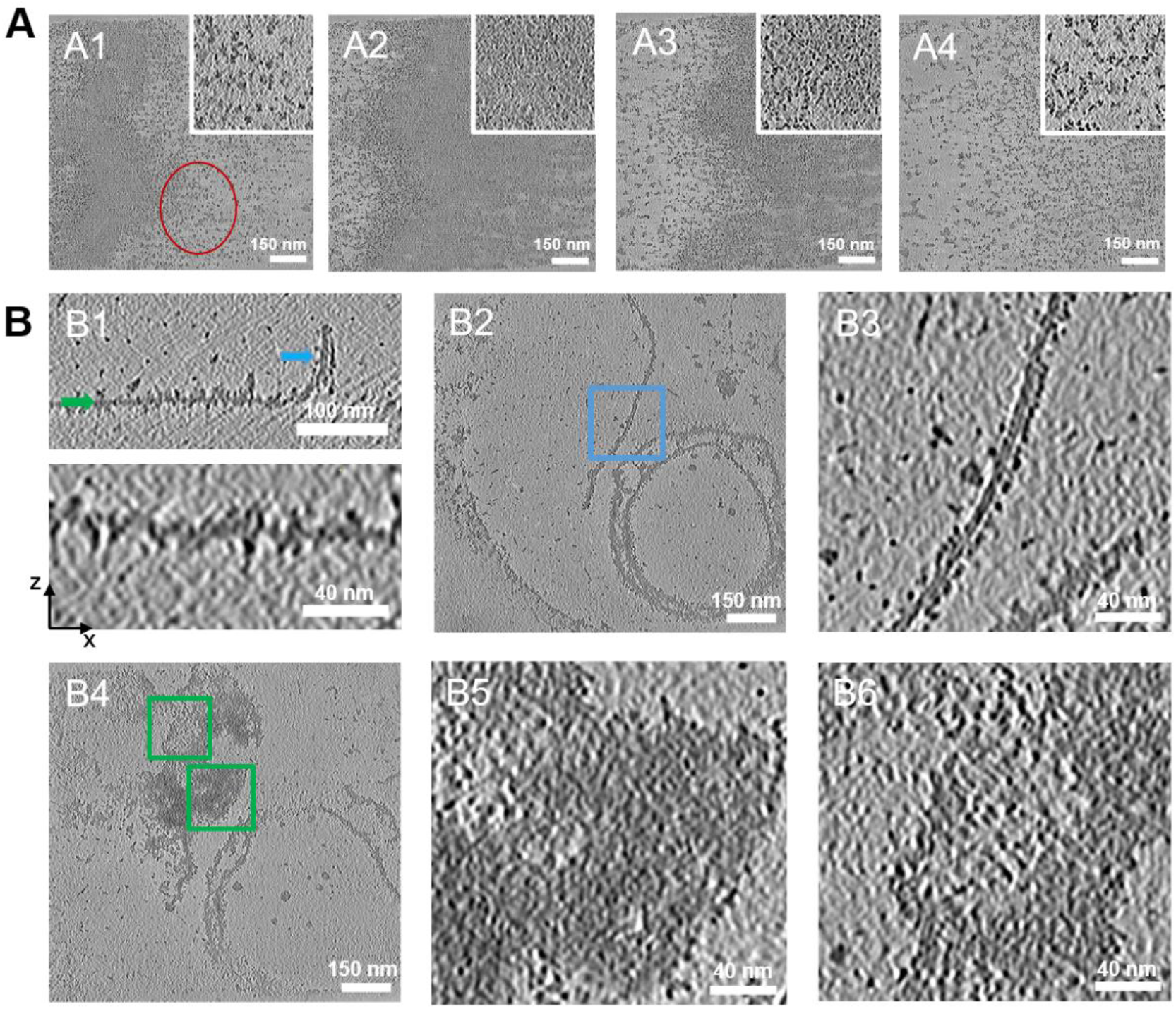
Overall morphology of plasma membranes. **(A)** Cryo-ET density slices from the 3D tomograms. A density slice of the outermost region of the ectoplasmic membrane **(A1)**, a very dense protein layer near the ectoplasmic membrane **(A2)**, a transmembrane region **(A3)** and a density slice on cytoplasmic side **(A4)** of the plasma membrane from the XY plane. Insets: enlargements of the same membrane region marked by a red circle. **(B)** Comparison of the characteristics of vertical and flat membranes. **(B1)** Cryo-ET density slices from the 3D tomograms of a curved membrane across the XZ plane, showing the existence of both vertical and flat sections on the membrane. The blue and green arrows indicate the corresponding Z-position at density slices shown in (B2) and (B4), respectively. A Cryo-ET density slice across the XY plane of the vertical section **(B2)** and flat section **(B4)** on a curved membrane. **(B3)** A magnified view of an area marked by a blue box in (B2) shows the appearance of a phospholipid bilayer and the transmembrane proteins across it. **(B5 and B6)** Magnified views of areas marked by green boxes in (B4), show the morphology of a transmembrane region and membrane proteins in a flat state.

Additionally, we collected some cryo-ET data from plasma membranes in a special state to further reveal each component in normal plasma membranes. For example, there were vertical and flat sections from density slices across the XZ plane of the 3D tomogram reconstructed from a curved 293T cell membrane (Fig. 2, B1). On the one hand, the lipid bilayer and the transmembrane proteins could be recognized distinctly from the vertical sections (Fig. 2, B2-B3). On the other hand, the size of transmembrane proteins and morphology of the transmembrane region could be observed conveniently from the flat sections of this curved 293T cell membrane (Fig. 2, B5-B6). Consequently, these observations provided supplementary evidence for improving the authenticity of the real plasma membrane structure we observed *in situ* by cryo-EM.

### Spatial distribution of membrane proteins and protein complexes

To further reveal the detailed information of membrane proteins and protein islands on the cytoplasmic side of plasma membranes, we quantitatively analysed the features of plasma membranes in the tomograms. First, the numbers of membrane proteins and protein islands on the cytoplasmic side of plasma membranes were counted. This particle concentration, measured as the number of membrane proteins and protein islands per μm^2^, ranged from ~8000 μm^-2^ to ~11000 μm^-2^ (Fig. 3, D). Second, 3D subareas of the cytoplasmic side of the plasma membrane were transformed into a 2D projection, and the area ratios of membrane proteins together with protein islands in the plasma membrane were calculated to be 25%-30% (Fig. 3, E), demonstrating the degree of compactness of membrane proteins on the inner side (*15*). Finally, the average sizes of all the particles on the cytoplasmic side of plasma membranes were measured to be approximately 7 nm in length and 4 nm in width (Fig. 3, F).

**Fig. 3.**
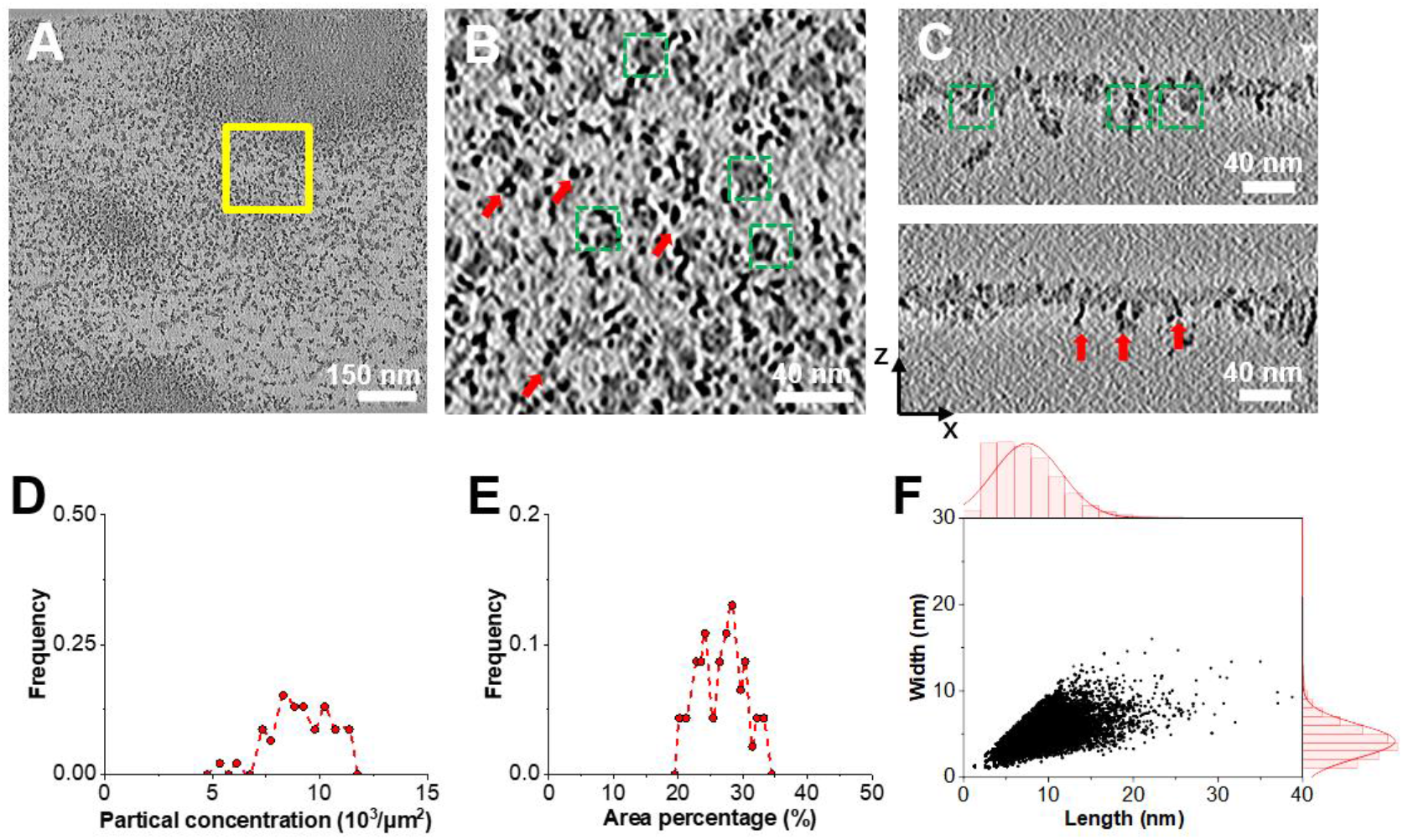
Statistical analysis of membrane proteins on the cytoplasmic side of plasma membranes. **(A)** A Cryo-ET density slice from the 3D reconstructed tomograms of the cytoplasmic side of a plasma membrane across the XY plane. **(B)** A magnified view of an area marked by a yellow box in (A). **(C)** Cryo-ET density slices across the XZ plane. The morphology of protein islands and their locations in a plasma membrane are marked out by green dashed boxes, and membrane proteins are pointed out by red arrows. Concentration **(D)**, area percentage **(E)**, length and width **(F)** of the membrane proteins and protein islands in the cytoplasmic side of the plasma membrane.

## Discussion

Structural biologists still have a long way to go before the *in situ* structure and detailed information of cell membrane are uncovered with nanometre resolution. In this study, we observed intact single-layer somatic cell membranes *in situ* with cryo-ET imaging for the first time and analysed the distribution of membrane proteins throughout whole regions of cell membranes.

Based on image processing of our cryo-ET data collected from single-layer 293T plasma membrane areas, we found that crowded membrane protein complexes together with ambient membrane proteins of similar heights (7-11 nm) exist on the ectoplasmic side of the plasma membrane, forming a dense protein layer on top of a lipid bilayer; proteins aggregate to form islands with different heights, up to tens of nanometres on the cytoplasmic side. Moreover, we determined the *in situ* distributions and relative locations of membrane proteins on the cytoplasmic side of the somatic plasma membranes by statistical measurement in the processed 3D tomograms.

It is worth noting that many protein–protein interactions take place on or near a cell membrane and result in the formation of protein clusters with different sizes, which we could observe from the 3D tomograms. We proposed three categories of clustering mechanisms that do not depend on the precise structure of membrane proteins: direct attractive interactions between membrane proteins facilitate clustering; the steric confinement of the cytoskeleton forces the proteins to become clusters; and the plasma membrane itself can generate forces between proteins(*16*). Further studies are needed to identify the molecular mechanisms and detailed structures of these membrane protein clusters.

To obtain clear and accurate cryo-ET data on somatic cell membranes, it is necessary for us to improve the membrane preparation method to obtain a cleaner cell membrane in an optimal native state for *in situ* cryo-ET imaging. Moreover, developments in instrumentation and image analysis will also help us to improve the resolution of our cryo-ET data. For example, we can acquire our cryo-ET data with a Volta phase plate (VPP) to enhance the contrast of the tomograms(*17*); and we can use IsoNet, a deep learning-based software package, to iteratively reconstruct the missing-wedge information and increase signal-to-noise ratio of the tomograms of our cryo-ET data(*18*); additionally, we can use Topaz-Denoise, a deep learning method which can learn the denoising process directly from our cryo-ET data, to denoise our cryo-EM images(*19*). We look forward to utilising these advanced approaches that facilitate the *in situ* imaging of somatic cell membranes by cryo-ET.

## Acknowledgements

We thank Z. Hong Zhou and Guo-Qiang Bi for technical advice on cryo-EM imaging and valuable suggestions on the manuscript. We thank the Cryo-EM Facility of Southern University of Science and Technology for providing technical support during EM image acquisition.

## Author contributions

C.L. performed the experiments. C.L. Y.Z. and T.Z. collected the data. Y.Z. processed and analysed the data. C.L. and Y.Z. wrote the manuscript. T.Z., G.Z. and HL.W. participated in data processing. H.W. conceived and designed the experiments, discussed the results and commented on the manuscript.

## Data and materials availability

All data are available in the main text.

